# Functional characterization of pathogenic SATB2 missense variants identifies distinct effects on chromatin binding and transcriptional activity

**DOI:** 10.1101/2025.05.28.656698

**Authors:** Joery den Hoed, Fleur Semmekrot, Jolijn Verseput, Alexander JM Dingemans, Dick Schijven, Clyde Francks, Yuri A. Zarate, Simon E. Fisher

## Abstract

SATB2-associated syndrome is an autosomal dominant neurodevelopmental syndrome caused by genetic alterations in the transcription factor SATB2. The associated phenotype is variable, and genotype-phenotype correlation studies have not yet been able to explain differences in severity and symptoms across affected individuals. While haploinsufficiency is the most often described disease mechanism, with the majority of variants consisting of whole- or partial-gene deletions and protein truncating variants with predicted loss-of-function, approximately one-third of affected individuals carry a *SATB2* missense variant with an unknown effect. In this study, we sought to functionally characterize these missense variants to uncover associated pathogenic mechanisms. We combined a set of human cell-based experiments to screen 31 etiological *SATB2* missense variants for effects on nuclear localization, global chromatin binding, and transcriptional activity. Our data indicate partial loss-of-function effects for most of the studied missense variants, but identify at least eight variants with increased SATB2 function showing a combination (or subset) of features that include stronger co-localization with DNA, decreased nuclear protein mobility suggesting increased overall chromatin binding, and maintained or increased transcriptional activity. These results demonstrate that phenotypes associated with variants in *SATB2* may have distinct underlying disease mechanisms, and the data could provide a resource for future studies investigating disease variability and potential therapies for this condition.

## Introduction

SATB2 is a transcription factor^1^ that is highly expressed during development in various organs and tissues, including the brain^2^, bones^3^ and intestines^4^. Heterozygous predominantly *de novo* variants in *SATB2* have been associated with an autosomal dominant neurodevelopmental syndrome, called SATB2-associated syndrome (SAS) or Glass syndrome (MIM 612313)^5; 6^. The phenotype of SAS is broad and highly variable across affected individuals, and includes developmental delay, absent or limited speech, craniofacial and dental problems, and skeletal anomalies as some of the core phenotypic features^7^.

To perform its function as a regulatory protein, SATB2 contains three DNA-binding domains: two highly homologous CUT domains (CUT1 and CUT2) and a homeobox domain (HOX; Figure 1)^1^. Prior investigations into four *SATB2* missense variants, located in the CUT domains, found that variants in CUT1 cause a decrease in SATB2 DNA-binding, while CUT2 missense variants have an opposite effect^8^. Based on these differences, distinct functions have been proposed for CUT1 and CUT2, with the CUT1 domain potentially being important for DNA binding and the CUT2 domain playing a role in disassociation from the DNA.

**Figure 1.**
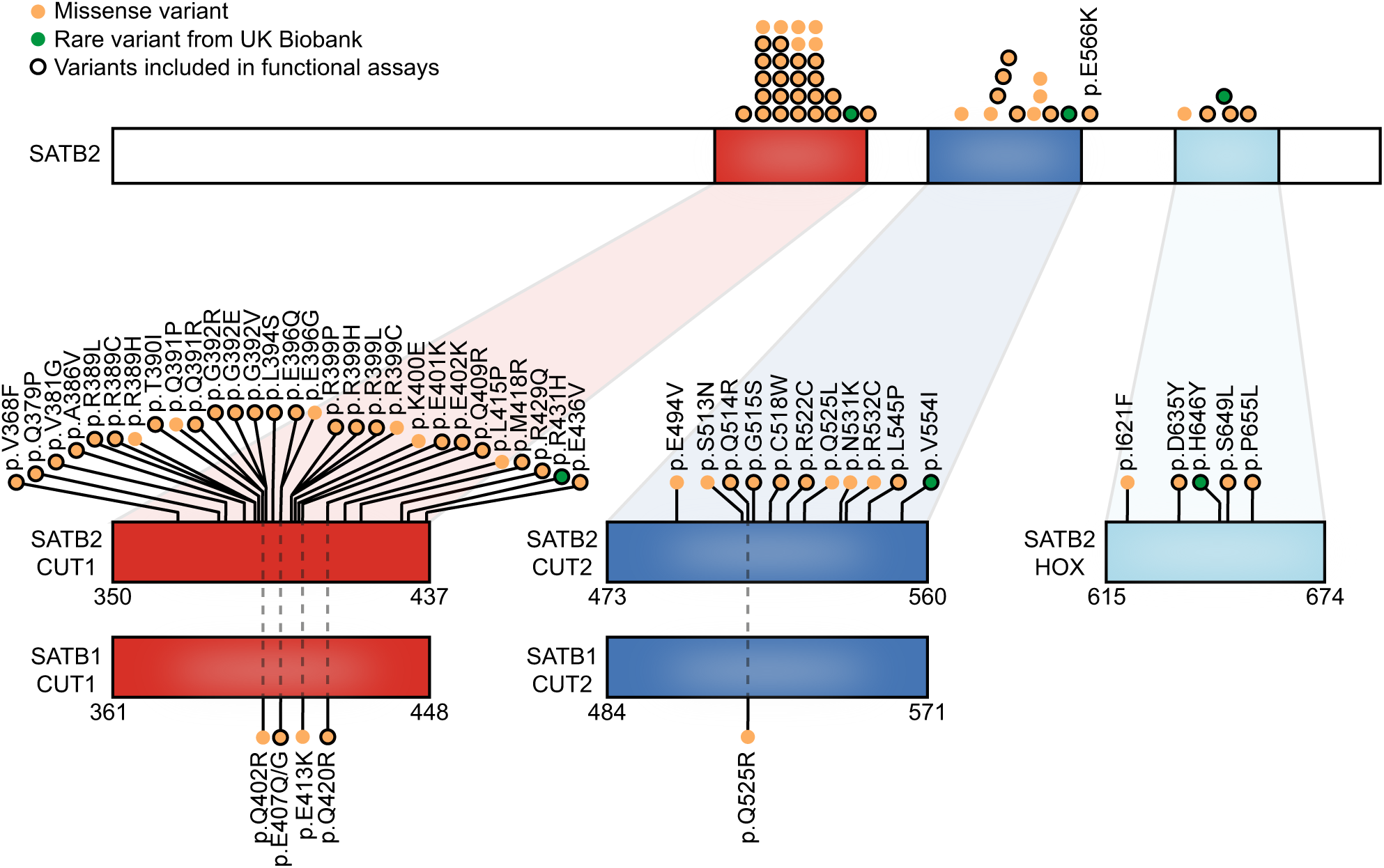
SATB2 missense variants located in the DNA binding domains and associated with neurodevelopmental disorder. Schematic representation of SATB2 (GenBank: NM_001172509.2/NP_001165980.1), including the DNA binding domains CUT1 (red), CUT2 (blue) and HOX (light blue). Etiological SATB2 missense variants present in the SATB2 Portal^10^ and located in the DNA-binding domains are shown in orange, and three selected rare missense variants from the UK Biobank are depicted in green. Black-encircled variants were included in functional assays. Below, the DNA binding domains of paralog SATB1 (GenBank: NM_001131010.4/NP_001124482.1) are shown with pathogenic SATB1 missense variants described in neurodevelopmental disorder affecting equivalent positions to etiological SATB2 variants.

Variants reported for SAS include whole- and partial-gene deletions, protein truncating variants, splice variants, and missense variants^5; 6^. Most of these etiological variants, identified in about 60-70% of affected individuals, have a predicted loss-of-function effect^6; 9^; therefore haploinsufficiency is the most widely proposed underlying mechanism of SAS. However, the remaining 30-40% of individuals with SAS are reported to carry a *SATB2* missense variant. These missense variants cluster in the three DNA-binding domains, with some variants showing high recurrence, although it remains unclear if they have loss-of-function effects as well^10^.

Multiple lines of evidence suggest that these missense variants may act differently from loss-of-function variants. In an earlier study proposing distinct functions for the CUT1 and CUT2 domains^8^, differences in functional consequences could have arisen from variant-specific rather than domain-specific effects. Moreover, pathogenic missense variants in the conserved CUT1 and CUT2 domains of the close family member SATB1, some of which affect residues equivalent to the ones affected by etiological *SATB2* variants (Figure 1), were recently described to cause stronger DNA-binding and increased transcriptional activity^9^. Lastly, we recently described three different amino acid changes affecting the same SATB2 CUT1 residue, p.Gly392, that showed distinct functional effects and resulted in significant differences in clinical outcome, specifically in the gross motor domain^11^. Together, these studies suggest that *SATB2* missense variants may make up a heterogeneous group with an array of differing functional consequences.

Genotype-phenotype correlations, based on comparisons between predicted loss-of-function and missense variants or on variant location, have only revealed subtle differences, being unable to explain phenotypic variability in SAS^6; 10^. Most prior functional work using cellular or animal models has investigated the effects of *SATB2* haploinsufficiency or complete loss of the gene^12–15^. To better understand phenotypic variability in SAS, studies are needed that map out the functional consequences of missense variants, in order to stratify the SAS population and to perform functionally-informed genotype-phenotype analyses.

In the present study, we used human cell-based assays to systematically screen the large majority of reported etiological *SATB2* missense variants in the DNA-binding domains (Figure 1)^6^ for their functional effects at the protein level. We found that while most CUT1 and CUT2 missense variants have a partial loss-of-function effect, a subset of variants shows increased SATB2 function. Our data demonstrate that etiological *SATB2* missense variants may underlie at least two, and potentially multiple, distinct disease mechanisms. Given that missense variants represent approximately one-third of causal variants in SAS, the functional data presented in this study will be crucial for performing more sophisticated genotype-phenotype analyses, and may eventually steer efforts to develop therapeutic strategies.

## Results

### Etiological SATB2 missense variants affect nuclear localization and co-localization with DNA

We performed functional analyses for 31 etiological *SATB2* variants located in the CUT1, CUT2, and HOX DNA binding domains, as well as three rare missense variants from the UK Biobank identified in healthy individuals with normal range fluid intelligence scores (p.Arg431His, p.Val5541Ile and p.His646Tyr; Figure 1, Figure S1). The SATB2 variants were expressed as N-terminal YFP-fusion proteins in HEK293T/17 cells. Overexpressed wild-type SATB2 and proteins carrying the rare UK Biobank missense variants localized mostly to the nucleus in a generally diffuse slightly granular pattern (Figure 2A, Figure S2). In contrast, for the large majority of etiological CUT1 and CUT2 missense variants there was significant aggregation of the protein, with the exception of p.Gln391Arg, p.Gly392Arg, and p.Leu545Pro (Figure 2A-D, Figure S2). The p.Cys518Trp variant showed a more diffuse nuclear localization pattern compared to the reference SATB2 protein. Interestingly, aggregation of proteins carrying etiological missense variants was not similarly distributed in the nucleus across all variants. For a subset of missense variants the protein formed a cage-like clustered nuclear pattern strongly co-localizing with the AT-rich DNA binding dye Hoechst 33342, comparable to what has been described for SATB1 missense variants associated with den Hoed-de Boer-Voisin syndrome (DHDBV, MIM 619229; SATB2 p.Glu396Gln, p.Glu401Lys, p.Glu402Lys, p.Gln409Arg, p.Glu436Val, p.Gln514Arg, p.Gly515Ser and p.Glu566Lys)^9^. The other variants that aggregated were typically clustered in the inner regions of the nucleus, showing an inverse localization pattern with Hoechst 33342 (Figure 2A-D, Figure S2). These results were replicated for a selection of CUT1 and CUT2 missense variants with the histone marker H2B, overexpressed as an RFP670nano3-fusion protein (Figure S3)^16^. A number of the variant proteins co-localizing with the DNA also showed a higher overall proportion of nuclear localization (p.Gln409Arg and p.Glu436Val), while some variant proteins with decreased co-localization with DNA showed reduced nuclear localization (p.Ala386Val, p.Arg389Leu, p.Thr390Ile, p.Gly392Glu and p.Arg429Gln; Figure S4). The three etiological HOX domain variants all caused formation of nuclear condensates, albeit different in distribution and size for each variant (Figure 2A), reminiscent of SATB1 protein truncating variants that disrupt the HOX domain^9^. These results suggest the existence of at least two functionally distinct subgroups of SATB2 missense variants in the CUT1 and CUT2 domains.

**Figure 2.**
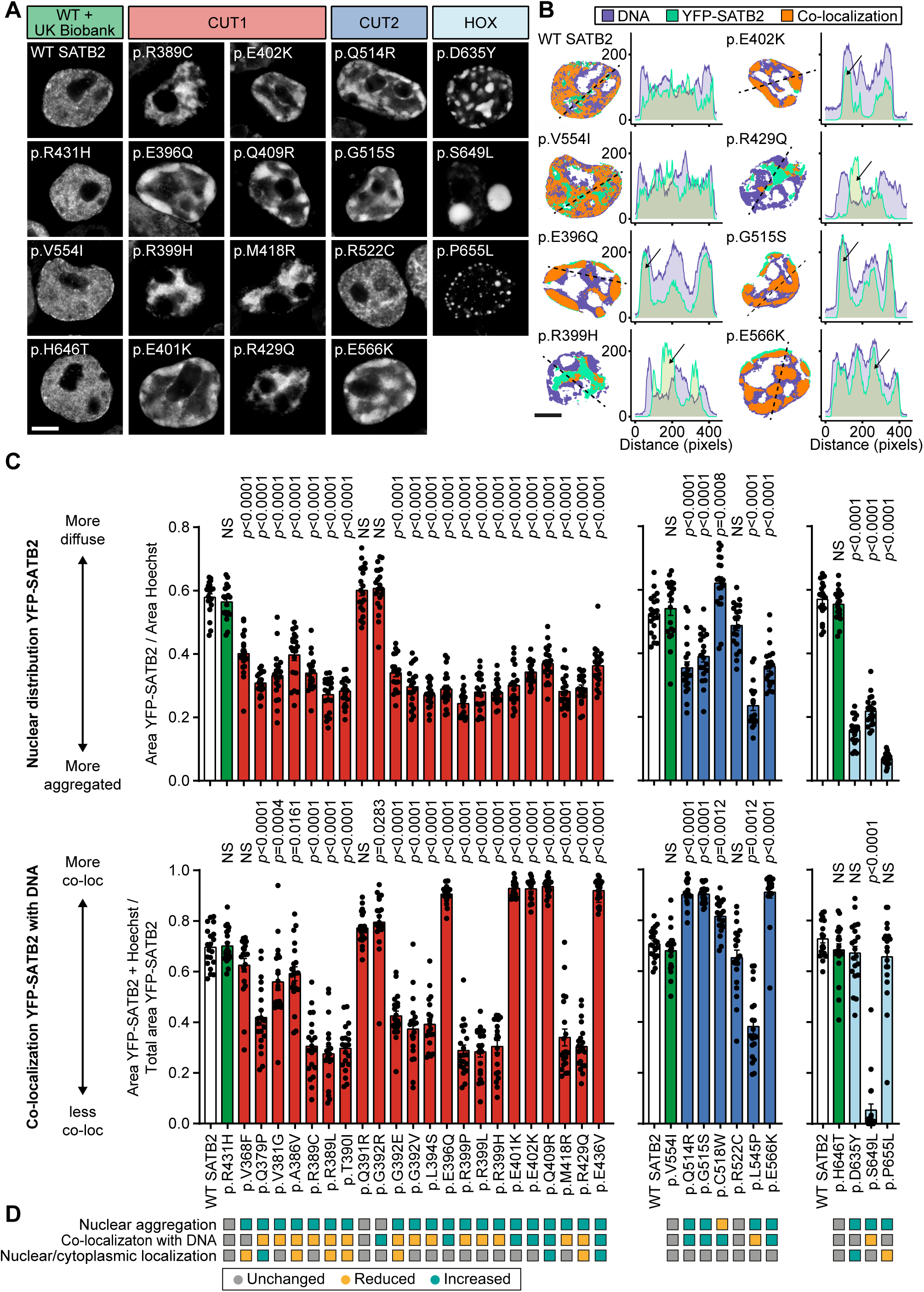
SATB2 missense variants affect nuclear localization. **A**) Direct fluorescence super-resolution imaging of nuclei of HEK293T/17 cells expressing YFP-SATB2 and variants. **B)** Left, thresholded masks of micrographs, showing the nuclear distribution of SATB2 (green) and Hoechst 33342 (magenta), and regions of co-localization (orange). Right, graphs depicting the intensity profiles of YFP-tagged SATB2 and variants, and the DNA binding dye Hoechst 33342. The profiles represent the fluorescence intensity values of the position of the dotted line shown in the threshold masks and drawn to cover both signals while avoiding the nucleoli (left). For each condition a representative image and corresponding intensity profile plot is shown. **A-B**) Panels show a selection of variants, other tested variants are included in Figure S2. Scale bar = 5 μm. **C**) Top, quantification of the nuclear distribution of YFP-SATB2. Bottom, quantification of co-localization of YFP-SATB2 with DNA binding dye Hoechst 33342. The UK Biobank variants are shaded in green, CUT1 domain variants in red, CUT2 domain variants in blue, and the homeobox variant in light blue. Values represent the mean ± SEM (*n* = 20 nuclei, *p* values compared to wildtype SATB2 [WT; white], one-way ANOVA and *post hoc* Dunnet test). **D**) Summary of image analysis results.

### Etiological SATB2 missense variants affect transcriptional activity

Next, we studied the effects of SATB2 missense variants on transcriptional activity using luciferase reporter assays. We used two previously established downstream targets of SATB2, the AT-rich matrix associated region (MAR) of mouse *Ctip2* (*Bcl11b*; chr12:107,914,809-107,915,262 in mm10)^2; 17^ and the MAR of mouse *Nr4a2* (chr2:57,107,317-57,107,710 in mm10)^17^. Both had a high identity (>90%) with the human regulatory sequences, and the reference SATB2 protein indeed showed transrepressive activity on these targets with a reduction of luciferase activity of ∼75% (Figure 3A, Figure S5). While SATB2 protein carrying the UK Biobank variants did not differ from the reference in these assays, the majority of the CUT1 and CUT2 missense variants, as well as the HOX variants, showed reduced repression of the targets, consistent with a partial loss-of-function (Figure 3A-B, Figure S5). In contrast, the missense variant proteins that showed stronger co-localization with the DNA dye Hoechst 33342 and H2B (Figure 2, Figure S2 and S3) demonstrated either retained or stronger transcriptional repression (Figure 3A-B, Figure S5), suggesting that these variants lead to stronger transcriptional repression of targets by SATB2 (p.Glu396Gln, p.Glu401Lys, p.Glu402Lys, p.Gln409Arg, p.Glu436Val, p.Gln514Arg, p.Gly515Ser and p.Glu566Lys).

**Figure 3.**
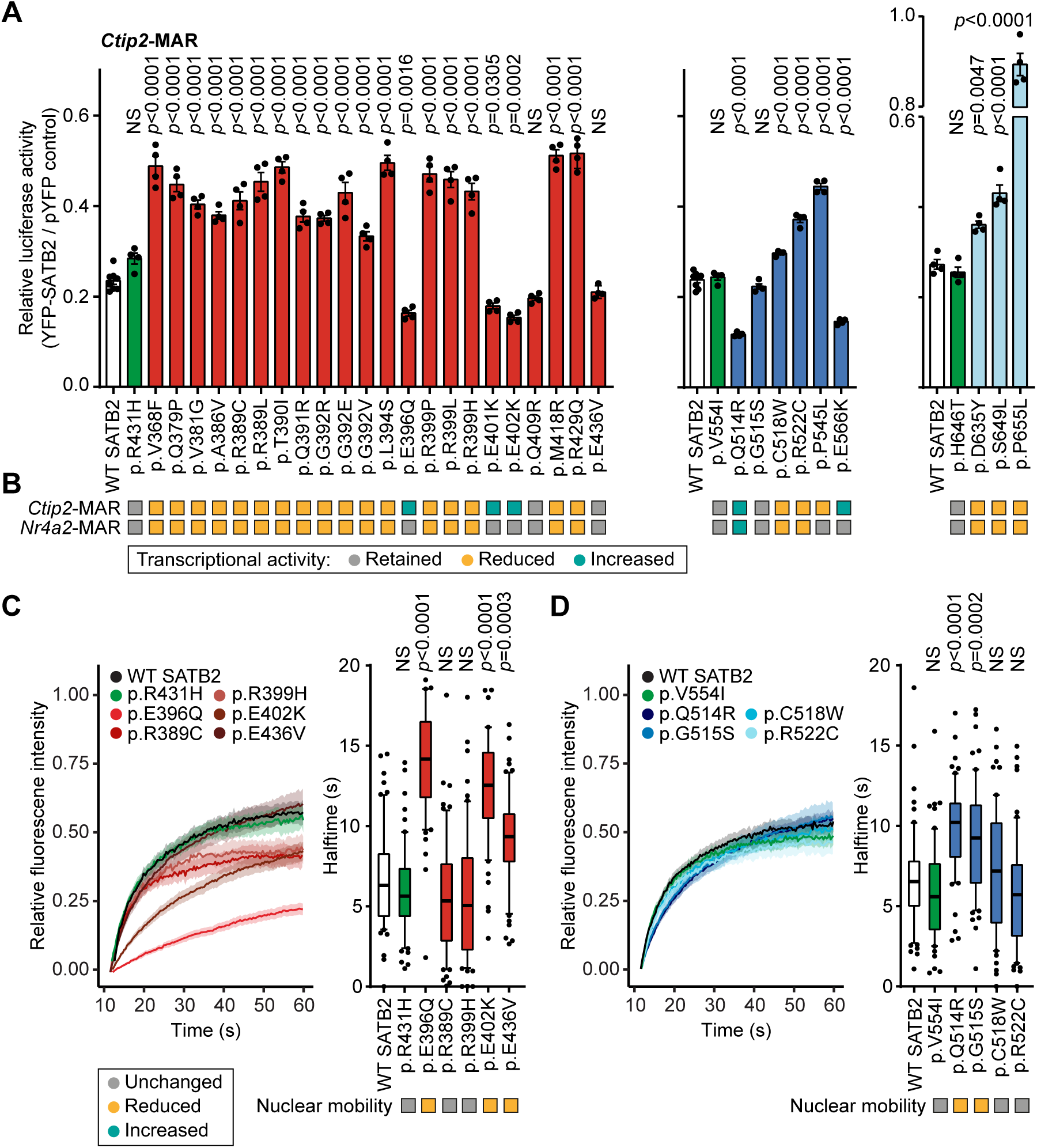
SATB2 missense variants affect DNA binding and transcriptional activity. **A**) Luciferase reporter assays using a reporter construct containing the mouse *Ctip2*-MAR4 binding site. UK Biobank variants are shaded in green, CUT1 domain variants in red, CUT2 domain variants in blue, and the homeobox variants in light blue. Values are expressed relative to the control (pYFP) and represent the mean ± SEM (*n* = 4-8, *p* values compared to wildtype SATB2 [WT; white], one-way ANOVA and *post hoc* Dunnet test). **B**) Summary of luciferase reporter assays of SATB2 transcriptional activity for both *Ctip2*-MAR and *Nr4a2*-MAR binding sites. Variants with a grey box retained their transrepressive activity, variants with a yellow box showed reduced repression and variants with a cyan-colored box increased repression. **B**) FRAP experiments to assess the dynamics of SATB2 CUT1 missense variants on chromatin binding in live cells. **D**) FRAP experiments to assess the dynamics of SATB2 CUT2 missense variants on chromatin binding in live cells. **C-D**) Left, mean recovery curves ± 95% CI recorded in HEK293T/17 cells expressing YFP-SATB2 fusion proteins. Right, box plots with median of the recovery halftime based on single-term exponential curve fitting of individual recordings (*n* = 60 nuclei from three independent experiments, *p* values compared to WT SATB2, one-way ANOVA and *post hoc* Dunnet test). Color code as in (**A**).

### Protein mobility assays suggest that SATB2 variants have distinct effects on transcription factor binding strength

To assess if these variants also affect the global dynamics of SATB2 chromatin binding, we selected a subset of variants for fluorescent recovery after photobleaching (FRAP) assays. We found that multiple variants (p.Glu396Gln, p.Glu402Lys, p.Glu436Val, p.Gln514Arg and p.Gly515Ser) had increased recovery half times, indicative of a reduced nuclear mobility and therefore stabilization of DNA binding, consistent with the luciferase reporter results (Figure 3C-D). In contrast, variants that showed reduced co-localization with DNA (Figure 2C) and/or transrepressive activity (Figure 3A-B) did not affect protein mobility in the nucleus (Figure 3C-D; p.Arg389Cys, p.Arg399His, p.Cys518Trp and p.Arg522Cys).

### Phenotypic analysis using facial photographs on functionally-informed subgroups

Next, we performed analysis using PhenoScore, an artificial intelligence-based phenomics framework that uses state-of-the-art facial recognition technology to quantify phenotypic similarity^18^. Comparing photographs of 148 individuals with different types of etiological *SATB2* variants (Figure S6A), to age-, sex- and ethnicity-matched controls with neurodevelopmental disorders, we found that individuals with *SATB2* variants have a significantly distinctive facial phenotype compared to the background of neurodevelopmental controls (AUC = 0.91, p value = 1.35×10^-6^). The predictive facial features were visualized in a LIME heatmap (Figure S6B). We continued by performing an analysis comparing photographs of 45 individuals with *SATB2* missense variants to matched individuals from the same SATB2 dataset with other variant types and found that individuals with missense variants have significant distinct facial features compared to individuals with *SATB2* variants other than missense variants (AUC = 0.66, p value = 0.022; Figure S6C). Finally, when comparing photographs of only individuals carrying a missense variant with a partial loss-of-function effect, as demonstrated in our functional assays, to photographs of matched individuals with a predicted full loss-of-function effect (Figure S6D), the facial features of these subgroups could not be significantly separated (AUC = 0.54, p value = 0.416). Too few photographs were available to perform any subgroup comparisons for *SATB2* missense variants causing an increase of SATB2 functions.

## Discussion

By screening 31 etiological *SATB2* missense variants using cell-based experiments (Figure 1, Figure 4), we show that missense variants associated with SAS may have distinct pathogenic mechanisms, with the majority showing effects consistent with a partial loss-of-function, a subset causing an increase in SATB2 function and a small set of variants not mapping to either of these two functional groups.

**Figure 4.**
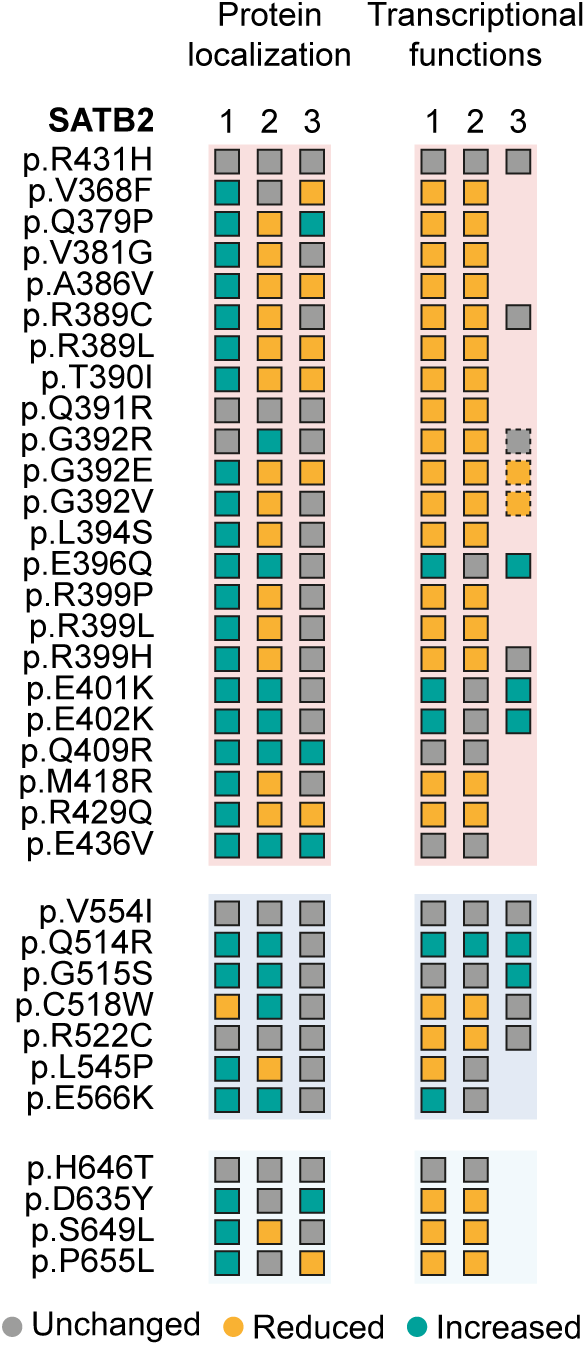
Summary of results for SATB2 missense variants across all functional read-outs assessed in this study. Overview of the functional effects of SATB2 variants in human cell-based assays. Protein localization (1) represents the results on nuclear aggregation, protein localization (2) on co-localization with DNA-dye Hoechst and histone marker H2B, and protein localization (3) the ratio of nuclear over cytoplasmic localization. Transcriptional functions (1) shows the results of the transcriptional reporter assays for the *Ctip2*-MAR binding site, transcriptional functions (2) the transcriptional reporter assays for the *Nr4a2*-MAR site, and transcriptional functions (3) the FRAP assays for global chromatin binding. The dashed squares represent results from FRAP assays performed by den Hoed et al. 2024 (Ref. 11).

Comparing our data to the limited number of prior functional investigations of missense variants (including p.Arg389Cys, p.Gly515Ser, and p.Glu566Lys), the effects on nuclear localization and protein mobility appear consistent^8^. In addition, functional studies of the homologous transcription factor SATB1 show striking similarities regarding the effects of missense variants equivalent between SATB1 and SATB2 (Figure 1; SATB2 Glu396Gln, p.Glu402Lys, p.Gln409Arg, p.Gln514Arg), also showing cage-like localization patterns for the variant proteins, as well as strong co-localization with DNA and increased transcriptional activity^9^. However, results from our cell-based assays are not in line with the earlier hypothesis on the distinct roles of the CUT1 and CUT2 domains controlling SATB2 association/disassociation with the chromatin^8^. While only one CUT1 variant (p.Arg389Cys), and two CUT2 variants were tested previously (p.Gly515Ser and p.Glu566Lys), either having a partial loss-of-function effect or causing increased chromatin binding respectively, our much larger functional screen shows that missense variants in both DNA-binding domains can have opposite effects on key SATB2 functions including DNA binding. Therefore, our data suggest that both CUT domains play a role in association with the DNA, consistent with various studies on these domains in SATB1 and SATB2^19^, and that the distinct functional effects on protein functions we observed in our assays are most likely variant-specific.

Moreover, our results replicate findings from prior investigations of the p.Gly392Arg, p.Gly392Glu and p.Gly392Val variants^11^, confirming that p.Gly392Arg behaves differently from the p.Gly392Glu and p.Gly392Val substitutions, with the latter two showing partial loss-of-function. In the cellular assays, the neighbouring variant, p.Gln391Arg, behaved very similarly to p.Gly392Arg, and the clinical phenotype associated with this variant matches the more severe phenotype described for p.Gly392Arg, including seizures, as well as non-ambulatory and non-verbal presentation at age five. These findings suggest that p.Gln391Arg potentially belongs to the same functional subgroup as p.Gly392Arg.

After our functional screen had already initiated, a novel *SATB2* missense variant in the CUT2 domain was reported: p.Glu519Lys^20^. The individual with this variant had a much more severe phenotype than generally observed for SAS, including lissencephaly. Interestingly, this variant affects the position within the CUT2 domain that is equivalent to the CUT1 p.Glu396Gln included in our assays (Figure 1), which showed increased SATB2 function. Also, the individual with the p.Glu396Gln variant was described with a particularly severe developmental phenotype with Rett-like features^21^. Moreover, the *SATB2* p.Glu519Lys variant is equivalent to *SATB1* CUT1 p.Glu530Lys, the most recurrent *SATB1* variant reported in DHDBV syndrome (8 out of 30 individuals reported with missense variants), which has been shown to cause increased DNA binding and transcriptional repression^9^. Therefore, it is very likely that *SATB2* p.Glu519Lys results in increased SATB2 function as well. As both p.Glu396Gln and p.Glu519Lys affect a core residue of the CUT domain^9^ and result in particularly severe phenotypes^20; 21^, they may be considerably detrimental to SATB2 protein function.

The number of missense variants that we identified to cause an increase in SATB2 functions make up a significant portion of the tested variant set (at least 8, 8/31 = ∼25%). However, the number of affected individuals with these variants represent only a small part of the overall SAS population (11 out of 435 total entries in the SATB2 Portal, 13/435 when the functionally uncharacterized p.Glu396Gly and p.Glu519Lys variants are included, and 11/165 when only missense variants are considered)^10^. Thus, in addition to the 60-70% of the SAS population carrying variants with a predicted loss-of-function effect, among individuals with pathogenic *SATB2* missense variants, partial loss-of-function is the main mechanism. The small number of individuals with missense variants that cause an increase in protein functions presented complications for our functionally-informed genotype-phenotype analyses, as the subgroup size of only five individuals with variants with increased SATB2 function was too small to perform facial photograph-based comparative analyses. The comparison of 24 individuals with partial loss-of-function *SATB2* missense variants to matched cases with predicted SATB2 complete loss-of-function did not indicate significant differences, suggesting that any facial differences may be very subtle. Larger phenotypic datasets will be needed to effectively perform such type of analyses in the future.

Even though the majority of the *SATB2* missense variants act as partial loss-of-function alleles, they may potentially still have distinct molecular effects from whole- and partial-gene deletions and protein-truncating variants with predicted full loss-of-function effects. SATB2 forms dimers or tetramers^22–24^ in order to perform its functions as a transcription factor. Moreover, SATB2 has been shown to be part of large chromatin remodeling complexes^25; 26^, steering them to target sites in the genome^25; 27^. Recently, etiological missense variants in the CUT1/2 domains of *SATB1*, some of which affect equivalent positions of *SATB2* missense variants included in this study, were shown to still form di/tetramers with wild-type SATB1^9^. Given the strong conservation between SATB1 and SATB2 (58% amino acid identity), this may be similar for *SATB2* missense variants as well, which suggests that *SATB2* missense variants may have potential dominant-negative effects, capturing away wild-type SATB2 through di/tetramerization and affecting larger protein complexes via intact protein-protein interactions.

Lastly, using our cell-based assays we investigated three missense variants in the HOX domain, one of which occurs recurrently in seven affected individuals listed in the SATB2 Portal^10^. Although the HOX variants showed a loss of transcriptional repression, consistent with a loss-of-function effect, they also caused abnormal aggregating localization patterns in the nuclei of cells. The nuclear punctae caused by HOX missense variants were different from what we observed for CUT1/2 missense variants, but similar to what has been seen before for synthetic^8; 9; 28^, and etiological^9^ SATB1 and SATB2 truncations that lack the HOX domain. The HOX domain has been shown to be crucial for recognition and binding affinity of transcriptional target sites for SATB transcription factors^19; 29^, but based on our (and prior) localization studies it may play a role in liquid-liquid phase separation of SATB2 as well, a function that would be interesting to further explore in future studies.

Taken together, using a set of cell-based functional assays, we show that etiological *SATB2* missense variants are a heterogeneous group that have a range of different functional consequences. Our data provide the first overview of the potential disease mechanisms associated with *SATB2* missense variants, show that haploinsufficiency is not the only underlying cause of SAS, and could be used as a guide for future molecular and clinical studies. Further characterization of the effects of *SATB2* variants will be important to continue refining studies that aim to understand phenotypic variability, which subsequently will be crucial for providing diagnostic care as well as steering therapeutic approaches.

## Methods

### UK Biobank variants for functional assays

For this study we used data from the UK Biobank^30; 31^, released in data release version 10.1 on the Research Analysis Platform (RAP) (https://ukbiobank.dnanexus.com). The UK Biobank received ethical approval from the National Research Ethics Service Committee North West-Haydock (reference 11/NW/0382), and all of their procedures were performed in accordance with the World Medical Association guidelines^32^. Written informed consent was provided by all of the enrolled participants. Some of our analyses were conducted on the UK Biobank RAP. We filtered the whole exome sequencing data (available for 454,707 individuals^31^) for rare missense variants located in the CUT1 (GRCh38 chr2:199328773-chr2:199348826), CUT2 (GRCh38 ch2:199308820-chr2:199323928) and HOX (GRCh38 chr2:199272391-chr2:199272570) domains of *SATB2* (ENST00000417098.6) and with an allele frequency of <0.1% (variants were annotated using snpEff v5.1d)^33^. We only included variants with more than one heterozygous carrier with a fluid intelligence score recorded on at least one assessment center visit (data-field IDs 20016 and 20191). Thirteen variants were identified, of which three (c.1936C>T [p.Arg431His], c.1660G>A [p.Val5541Ile] and c.1292G>A [p.His646Tyr]) were included in cell-based functional assays, one from each functional domain and located in close proximity to etiological *SATB2* missense variants. For these variants, fluid intelligence scores for respectively three, nine and two heterozygous individuals were available (Figure S1).

### Plasmids and site-directed mutagenesis

The cloning of SATB2 (NM_001172509) has been described previously^34^. Variants in SATB2 were generated using the QuikChange Lightning Site-Directed Mutagenesis Kit (Agilent) with primers listed in Table S1. cDNAs were subcloned using the *BclI*/*XbaI* restriction sites into pYFP, created by modification of the pEGFP-C2 vector (Clontech) as described before^35^. The pH2B-miRFP670nano3 construct^16^ was obtained from Addgene (plasmid #184670). Luciferase reporter plasmids pGL4.10-Ctip2-MAR4 and pGL4.10-Nr4a2-MAR4^17^ were gifted by Dr. Lei Zhang and Dr. Yu-Qiang Ding. All constructs were verified by whole-plasmid sequencing.

### Cell culture

HEK293T/17 cells (CRL-11268, ATCC) were cultured in DMEM supplemented with 10% fetal bovine serum and 1x penicillin-streptomycin (all Gibco) at 37 °C with 5% CO_2_. Transfections for luciferase reporter and FRAP assays were performed using GeneJuice (Millipore) following the manufacturer’s protocol. Transfections for immunoblotting and direct fluorescence microscopy were done with polyethylenimine (PEI; Sigma).

### Direct fluorescence microscopy

HEK293T/17 cells were grown on coverslips coated with poly-D-lysine (Sigma) and transfected with YFP-tagged SATB2 variants and RFP670nano3-tagged H2B. Cells were fixed with 4% paraformaldehyde (PFA, Electron Microscopy Sciences) 48 h after transfection, and nuclei were stained with Hoechst 33342 (Invitrogen). Fluorescence images were acquired with a Zeiss LSM880 confocal microscope and Airyscan unit with an alpha Plan-Apochromat 100×/1.46 Oil DIC M27 objective (all Zeiss) using a 4.5 zoom factor. Twenty images of single nuclei were taken for each variant. The raw CZI-format files were loaded into Fiji/ImageJ for further analyses. For each variant, the intensity profiles of the YFP-SATB2 and Hoechst 33342 or H2B-RFP670nano3 signal of a representative image were plotted using the ‘Plot Profile’ tool. The signals were also thresholded using the ‘Moments’ threshold to create representative SATB2/Hoechst 33342 or SATB2/H2B overlay images from the binary output. For quantification of aggregation, the YFP-SATB2 signal was thresholded with the ‘Moments’, and the Hoechst 33342 signal with the ‘Minimum’ threshold, and the areas of the binary thresholded images were measured. With these values a ratio was calculated: [Area thresholded YFP-SATB2 signal] / [Area thresholded Hoechst 33342 signal]. For quantification of co-localization of YFP-SATB2 with Hoechst 33342, both the YFP-SATB2 and Hoechst 33342 signals were thresholded with the ‘Moments’ threshold. Then, from the binary output a new image was created containing only pixels positive for both YFP-SATB2 and Hoechst 33342 using the ‘Image calculator’ tool. Areas of the binary thresholded images were measured and a ratio was calculated: [Area of pixels positive for both YFP-SATB2 and Hoechst 33342] / [Area thresholded YFP-SATB2 signal]. To determine the proportion of nuclear localization of YFP-SATB2 variants, ten regions of interest of 1000×1000 μm were imaged using a Zeiss AxioScan Z1 microscope. Raw CZI-files were loaded into Fiji/ImageJ. The YFP-SATB2 signal was thesholded with the ‘Li’ and the Hoechst 33342 signal with the ‘Otsu’ threshold. From the binary output a new image was created containing only pixels positive for both YFP-SATB2 and Hoechst 33342 using the ‘Image calculator’ tool. Areas of the binary thresholded images were measured and a ratio was calculated to determine the amount of nuclear YFP-SATB2 signal: [Area of pixels positive for both YFP-SATB2 and Hoechst 33342] / [Area thresholded YFP-SATB2 signal].

### Luciferase reporter assays

Luciferase reporter assays were performed with the pGL4.10-Ctip2-MAR4 and pGL4.10-Nr4a2-MAR4 reporter plasmids^17^. HEK293T/17 cells were transfected with one of the firefly luciferase reporters and a *Renilla* luciferase normalization control (pGL4.74; Promega) in a ratio of 50:1, as well as with YFP-SATB2 (WT or variant) or an empty control vector (pYFP). Forty-eight hours post-transfection, firefly and *Renilla* luciferase activity were measured using the Dual-Luciferase Reporter Assay system (Promega) with the Infinite M Plex Microplate reader (Tecan). After normalization for *Renilla* luciferase activity the data points were presented relative to the empty control vector condition.

### FRAP assays

FRAP assays were performed as described previously^9^. HEK293T/17 cells were cultured in clear-bottomed black 96-well plates and transfected with YFP-tagged SATB2 variants. Forty-eight hours after transfection, the culture medium was replaced with phenol red-free DMEM, containing 10% fetal bovine serum and 1x penicillin-streptomycin (all Gibco). The cells were then moved to a temperature-controlled incubation chamber at 37 °C. Fluorescent recordings were acquired using a Zeiss LSM880 confocal microscope and the Zen Black Image Software, with an alpha Plan-Apochromat 100×/1.46 Oil DIC M27 objective (Zeiss). FRAP experiments were performed by photobleaching an area of 0.98 μm x 0.98 μm within a single nucleus with 488-nm light at 100% laser power for 15 iterations with a pixel dwell time of 32.97 μm. The bleaching event was followed by collection of times series of 145 images with a 2.5 zoom factor and an optical thickness of 1.4 μm (2.0 Airy Units). Individual recovery curves were background subtracted and normalized for a bleach control. Then, the pre-bleach values were set at 100%, while the bleach value was set at 0%. The mean recovery curves were calculated using the EasyFRAP software^36^. Curve fitting was done with the Frapbot application using averaged normalization and a single-component exponential model^37^ to calculate the half-time of the recovery.

### Statistical analyses of cell-based functional assays

Statistical analyses for cell-based functional assays were carried out using a one-way ANOVA followed by a Dunnet *post hoc* test using GraphPad Prism Software. Statistical analyses for FRAP data were performed on values derived from fitted curves of individual recordings.

### Phenotypic subgroup analysis using facial photographs

Genotype-phenotype analysis on facial photographs was performed using PhenoScore^18^. From the 158 available photographs from individuals with SAS with all types of *SATB2* variants (derived from the SATB2 Portal^10^; Figure S6), 148 had sufficient quality and could be paired with a sex-, age- and ethnicity-matched control with a neurodevelopmental disorder from an in-house database from the Radboudumc. A comparison between photographs of individuals with SAS and with other neurodevelopmental disorders was performed as described previously^18^. Based on that comparison, a heatmap was generated using LIME, highlighting which facial areas were most important according to the established model. Subsequently, subgroup analyses within the SATB2 cohort were performed. The facial photographs of 45 individuals with missense variants were compared to the photographs of sex-, age- and ethinicity-matched individuals with other types of *SATB2* variants. Next, the facial photographs of individuals with SATB2 missense variants with a functionally-demonstrated partial LoF effect were compared to the photographs of individuals with variants with a predicted complete LoF effect (protein truncating variants and whole- or partial-gene deletions). After sex-, age- and ethnicity-matching across subgroups, 24 individuals could be included in the analysis. For a subgroup comparison for individuals with SATB2 missense variants that were found to cause an increase in SATB2 functions, five photographs were available, for which only four could be sex-, age- and ethnicity-matched with the other SATB2 subgroups, making the group size too small to perform a meaningful PhenoScore analysis, as was demontstrated in a prior study^18^.

## Acknowledgements

This work was financially supported by a research grant from the *SATB2* Gene Foundation and institutional funds from the Max Planck Society (Dr. S.E. Fisher and Dr. J. den Hoed). This research made use of data from the UK Biobank resource under Application Number 16066, with Dr. C. Francks as the principal applicant. We are grateful to all families participating in this study. Several authors of this publication (J. Verseput and Dr. A. Dingemans) are members of the European Reference Network on Rare Congenital Malformations and Rare Intellectual Disability ERN-ITHACA.

