## Supplemental Figures and Table for "Functional characterization of pathogenic SATB2 missense variants identifies distinct effects on chromatin binding and transcriptional activity"

| cDNA (ENSG00000119042) | Protein (ENST00000417098.6) | UK Biobank count and frequency | gnomAD (v2.1.1) count and frequency | Domain |
|---|---|---|---|---|
| c.1936C>T | p.Arg431His | 2 / $2.20 \times 10^{-6}$ | 1 / $3.98 \times 10^{-6}$ | CUT1 |
| c.1660G>A | p.Val554Ile | 17 / $1.87 \times 10^{-5}$ | 2 / $7.96 \times 10^{-6}$ | CUT2 |
| c.1292G>A | p.His646Tyr | 4 / $4.4 \times 10^{-6}$ | 2 / $7.09 \times 10^{-6}$ | HOX |

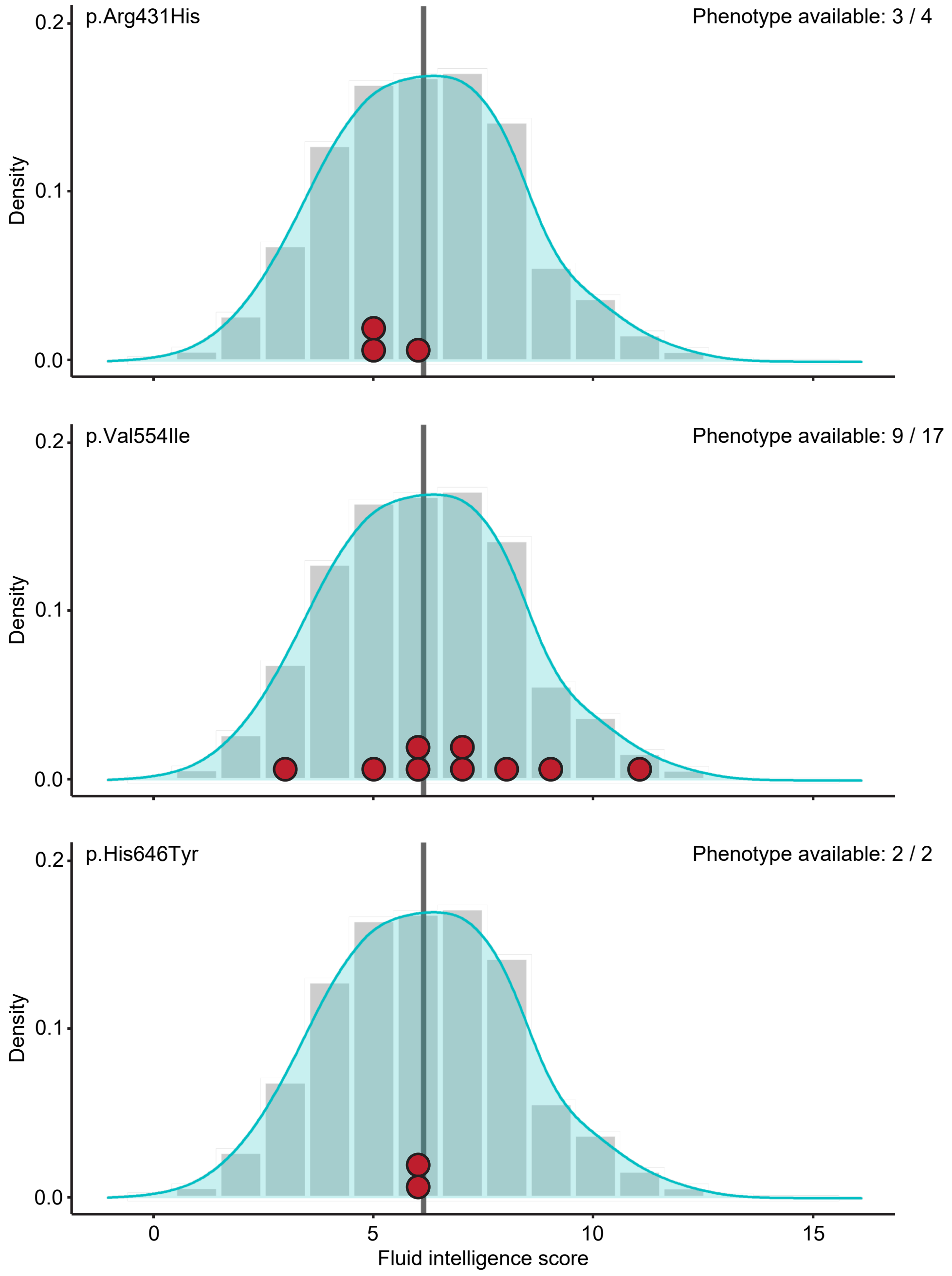


**Figure S1. Fluid intelligence scores of individuals from the UK Biobank with rare *SATB2* missense variants.** Top, a table with variant details and the frequencies of three rare *SATB2* variants from the UK Biobank that were included in the functional assays. Below, phenotype values of fluid intelligence of individuals with the selected rare *SATB2* variants in red, projected on the phenotype distributions of non-carriers (grey bars and blue shading).

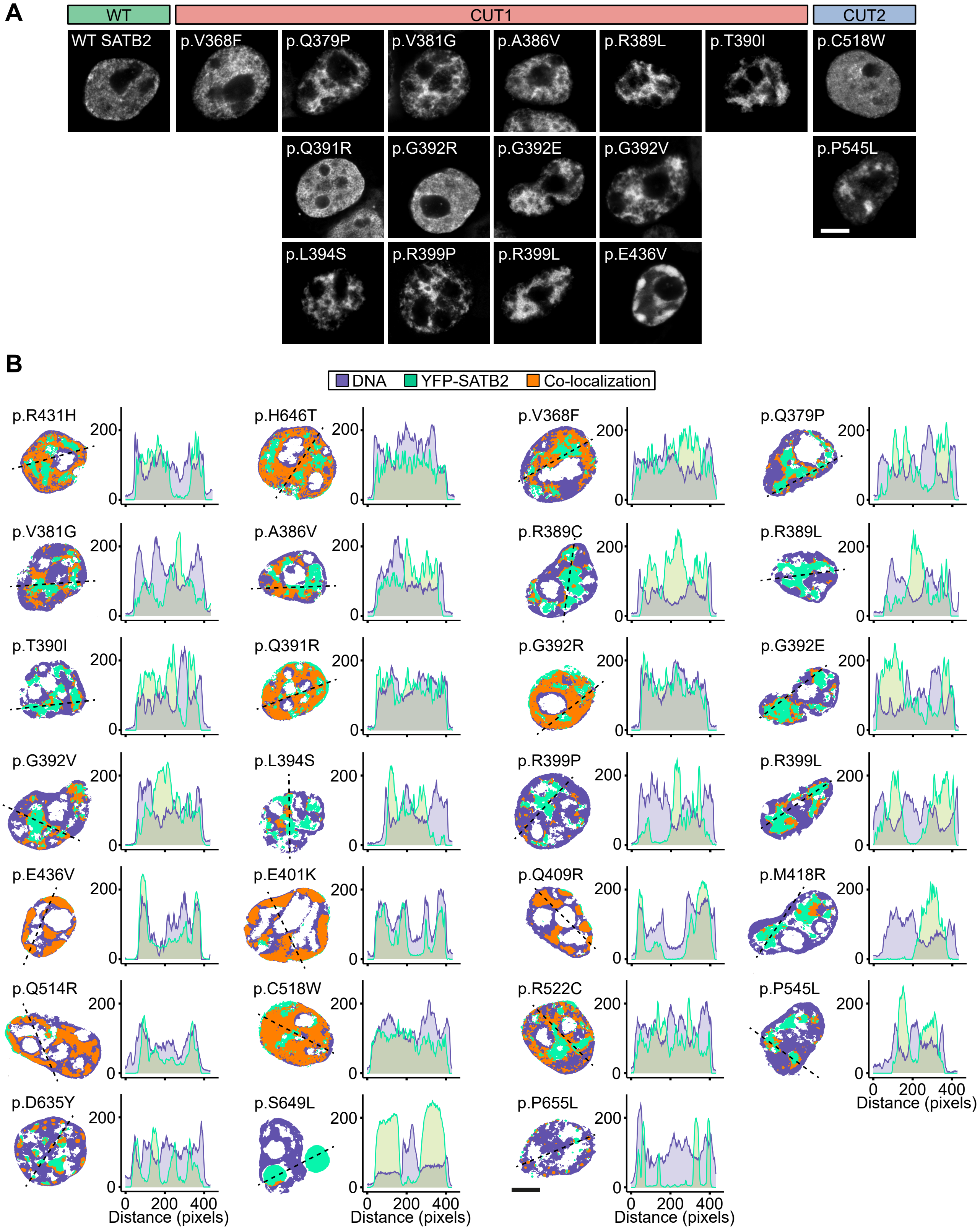


**Figure S2. SATB2 missense variants affect co-localization with the DNA. A**) Direct fluorescence super-resolution imaging of nuclei of HEK293T/17 cells expressing YFP-SATB2 and variants. **B**) Left, thresholded masks of micrographs, showing the nuclear distribution of SATB2 (green) and Hoechst 33342 (magenta), and regions of co-localization (orange). Right, graphs depicting the intensity profiles of YFP-tagged SATB2 and variants, and the DNA binding dye Hoechst 33342. The profiles represent the fluorescence intensity values of the position of the dotted line shown in the threshold masks and drawn to cover both signals while avoiding the nucleoli (left). For each condition a representative image and corresponding intensity profile plot is shown. **A-B**) Scale bar = 5 µm.

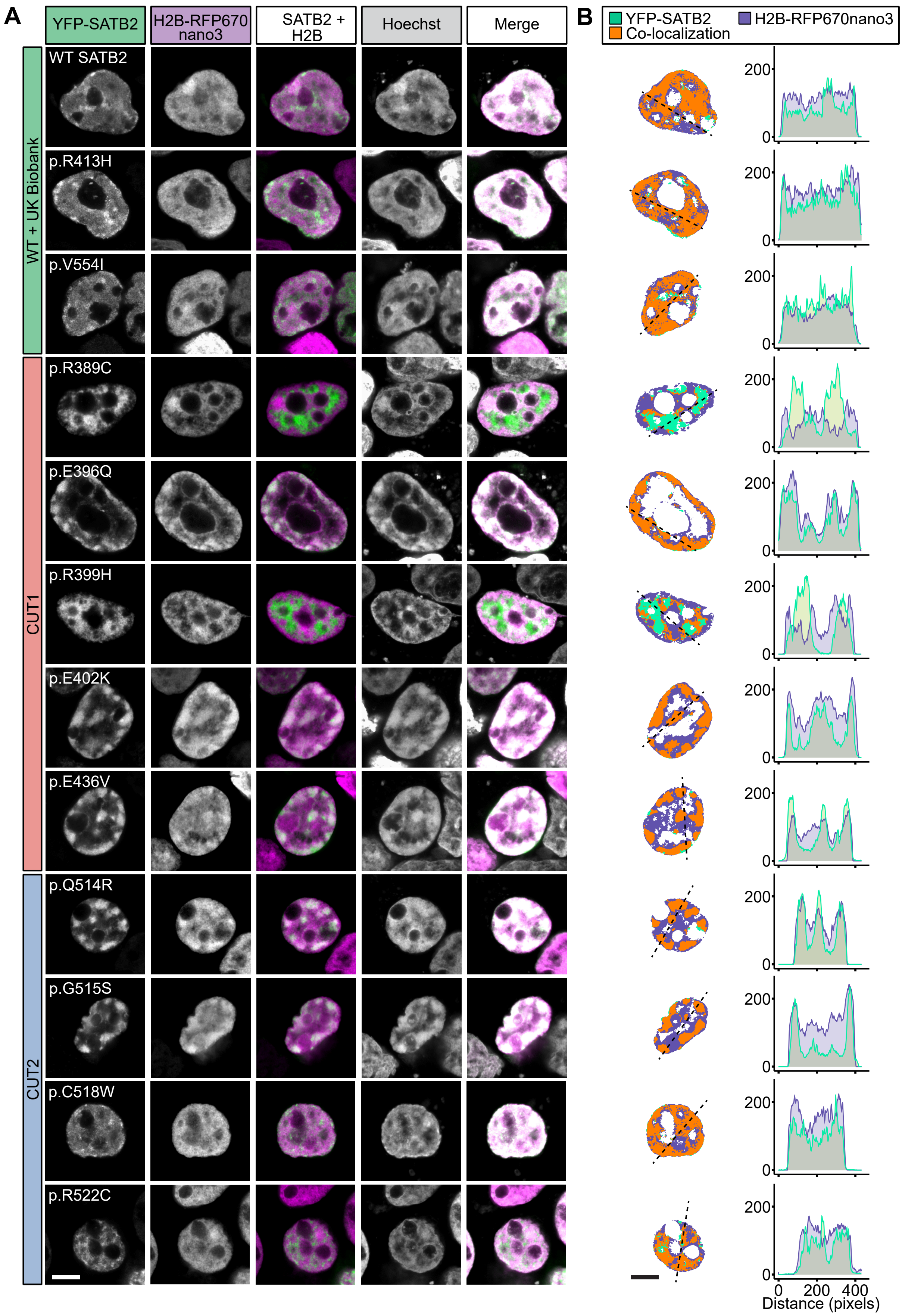


**Figure S3. SATB2 missense variants affect co-localization with the DNA. A**) Direct fluorescence super-resolution imaging of nuclei of HEK293T/17 cells expressing YFP-SATB2 and H2B-RFP670nano3 fusion proteins. SATB2 proteins are shown in green, histone marker H2B in magenta and the DNA binding dye Hoechst 33342 in white. **B**) Left, thresholded masks of micrographs, showing the nuclear distribution of SATB2 (green) and H2B (magenta), and regions of co-localization (orange). Right, graphs depicting the intensity profiles of YFP-tagged SATB2 and variants, and the histone marker H2B. The profiles represent the fluorescence intensity values of the position of the dotted line shown in the threshold masks and drawn to cover both signals while avoiding the nucleoli (left). For each condition a representative image and corresponding intensity profile plot is shown. **A-B**) Scale bar = 5 μm.

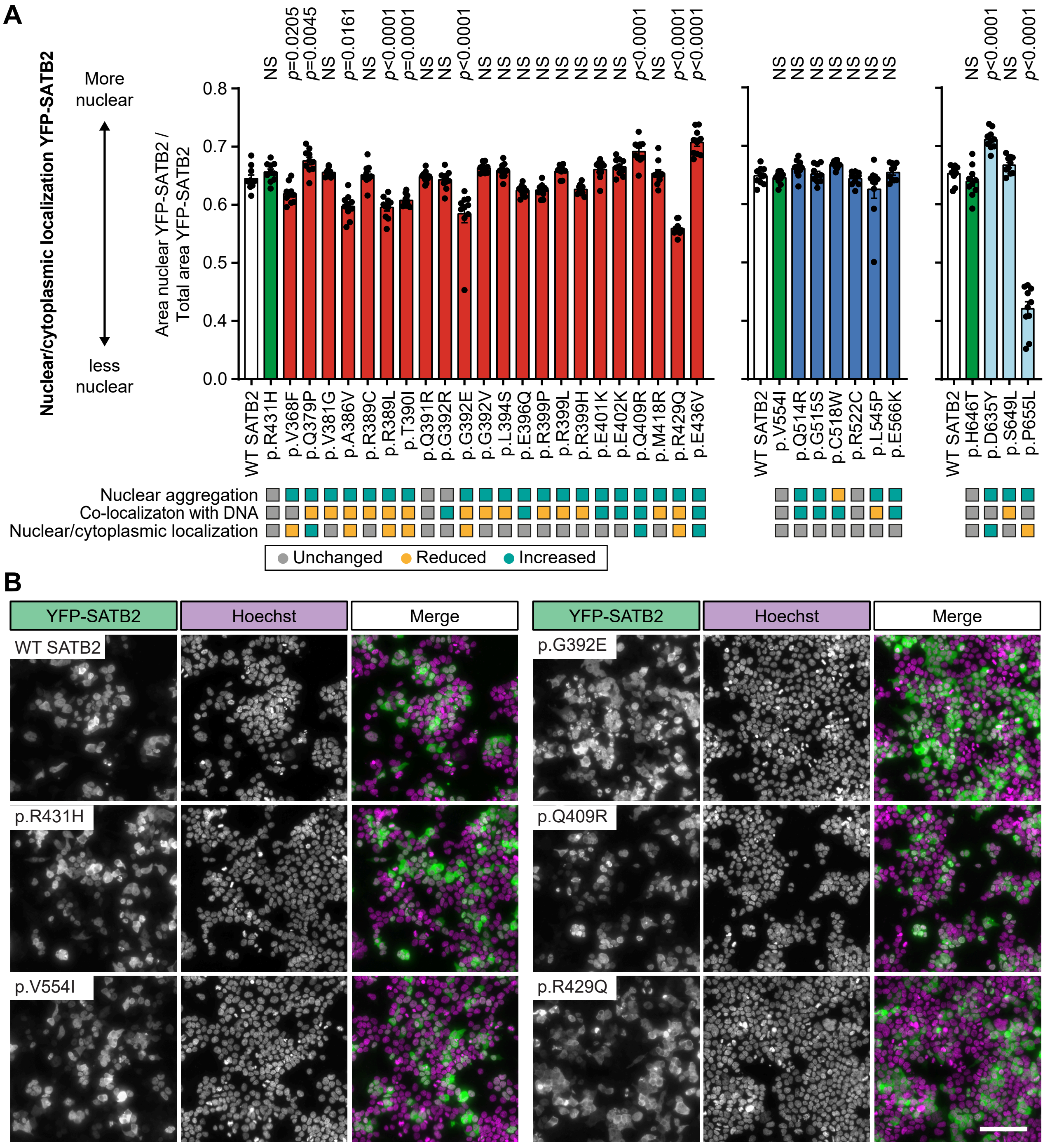


**Figure S4. A subset of SATB2 missense variants affect the nuclear to cytoplasmic localization ratio of the protein. A**) Quantification of the proportion of YFP-SATB2 signal localizing to the nucleus over the total signal of YFP-SATB2. The UK Biobank variants are shaded in green, CUT1 domain variants in red, CUT2 domain variants in blue, and the homeobox variant in light blue. Values represent the mean ± SEM (*n* = 10 regions of 1000 x 1000 µm, *p* values compared to wildtype SATB2 [WT; white], one-way ANOVA and post hoc Dunnet test). **B**) Representative micrographs of HEK293T/17 cells expressing YFP-SATB2 and a selection of variants to show the distrubtion of YFP-SATB2 in cells. SATB2 proteins are shown in green and the DNA binding dye Hoechst 33342 in magenta. Scale bar = 100 µm.

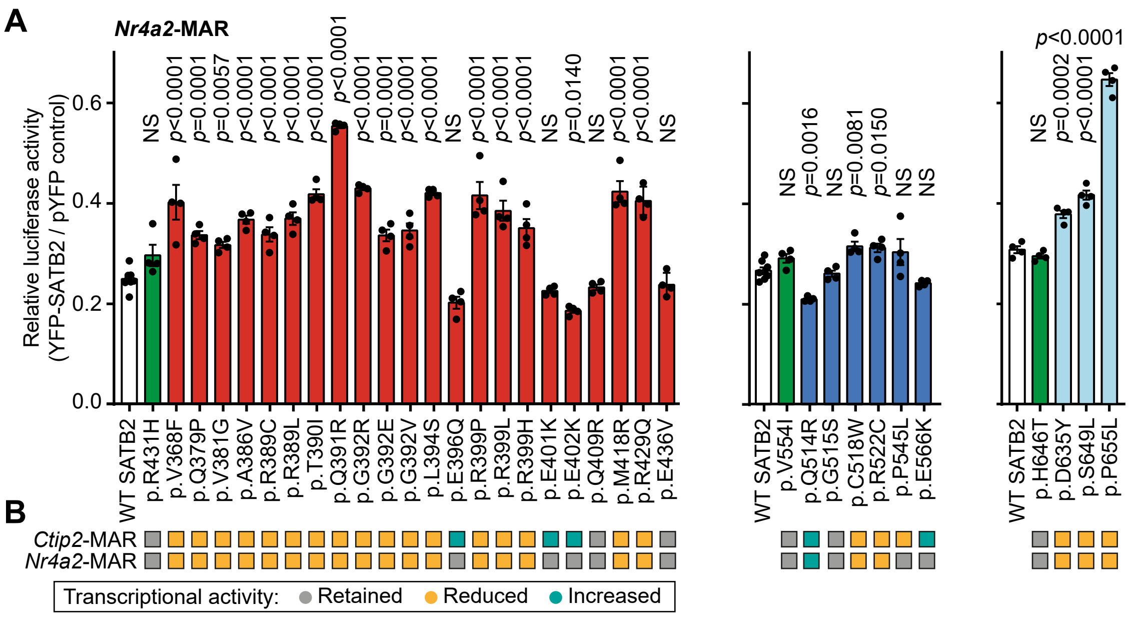


**Figure S5. SATB2 missense variants affect transcriptional activity. A**) Luciferase reporter assays using a reporter construct containing the mouse *Nr4a2*-MAR4 binding site. UK Biobank variants are shaded in green, CUT1 domain variants in red, CUT2 domain variants in blue, and the homeobox variants in light blue. Values are expressed relative to the control (pYFP) and represent the mean ± SEM ($n$ = 4-8, $p$ values compared to wildtype SATB2 [WT; white], one-way ANOVA and post hoc Dunnet test). **B**) Summary of luciferase reporter assays of SATB2 transcriptional activity for both *Ctip2*-MAR and *Nr4a2*-MAR binding sites. Variants with a grey box retained their transrepressive activity, variants with a yellow box showed reduced repression and variants with a cyan-colored box increased repression.

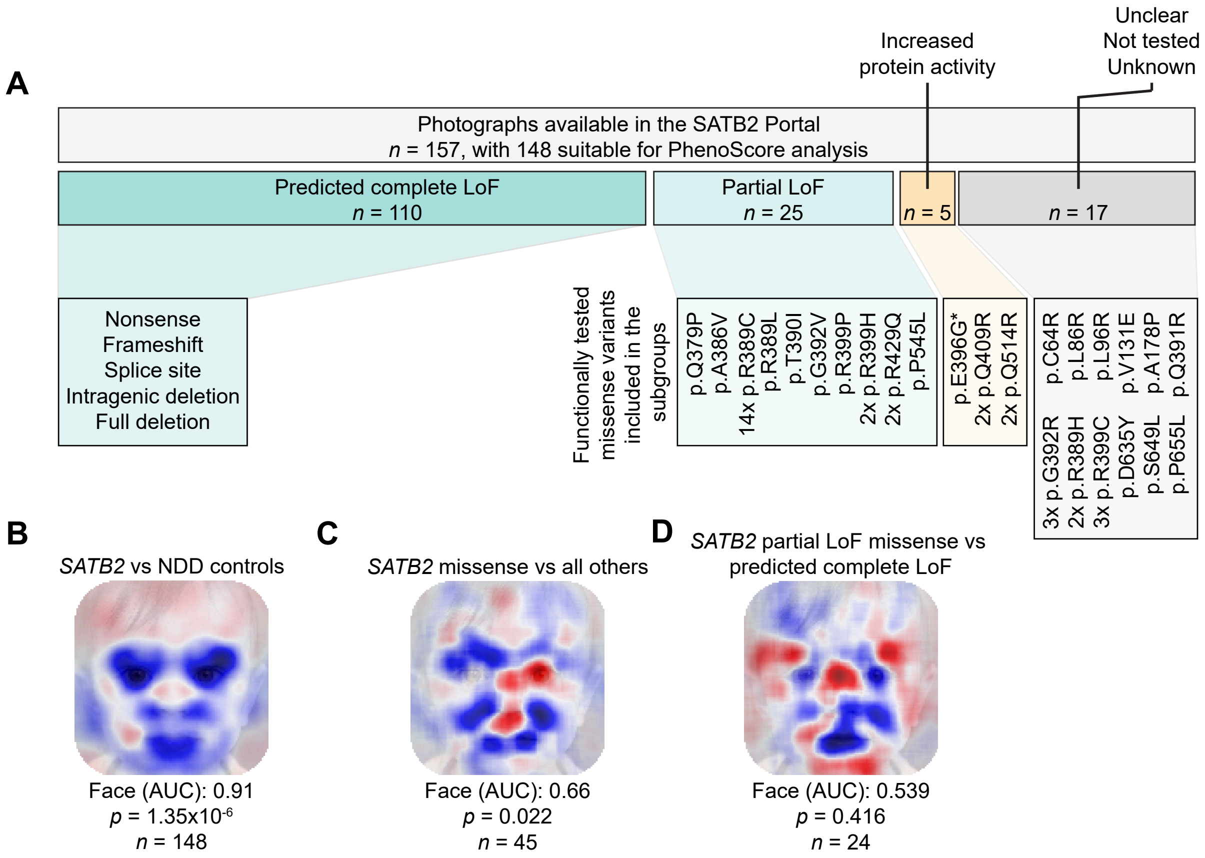


**Figure S6. Functionally-informed genotype-phenotype analysis on facial photographs using PhenoScore. A**) Overview of the available facial photographs in the SATB2 Portal (*n* = 157), with below the number of photographs for each functional subgroup, including variant details. **B**) A LIME heatmap indicating the facial areas most characteristic across individuals with *SATB2* variants when compared to controls with neurodevelopmental disorders (NDD). From the total of 157 photographs, 148 had a sex-, age- and ethnicity-matched NDD control, and could be included in the analysis. **C**) A LIME heatmap indicating the facial areas characteristic across all individuals with a *SATB2* missense variant compared to individuals with other *SATB2* variants. From the total of 47 individuals with a missense variant, 45 had a sex-, age- and ethnicity-matched control in the comparison group and could be included in the analysis. **D**) A LIME heatmap indicating the facial areas characteristic across individuals with *SATB2* missense variants with a partial loss-of-function (LoF) effect based on the cell-based functional assays, compared to individuals with a *SATB2* variant with a predicted complete LoF effect. From the 25 individuals with a partial LoF missense variant, 24 had a sex- and age-matched control in the comparison group and could be included in the analysis.

**Table S1. Primer sequences for site-directed mutagenesis**

| Primer | Primer Sequence (5' to 3') |
|---|---|
| SATB2-V368F-F | TCCAGATATCTACCAGCAATTCAGAGATGAGCTGAAG |
| SATB2-V368F-R | CTTCAGCTCATCTCTGAATTGCTGGTAGATATCTGGA |
| SATB2-Q379P-F | GGGCCAGTGTGTCCCCAGCTGTCTTTGCAAG |
| SATB2-Q379P-R | CTTGCAAAGACAGCTGGGGACACACTGGCCC |
| SATB2-V381G-F | GTGTGTCCCAAGCTGGCTTTGCAAGAGTGGC |
| SATB2-V381G-R | GCCACTCTTGCAAAGCCAGCTTGGGACACAC |
| SATB2-A386V-F | TGTCTTTGCAAGAGTGGTATTCAACCGCACACAGG |
| SATB2-A386V-R | CCTGTGTGCGGTTGAATACCACTCTTGCAAAGACA |
| SATB2-R389C-F | CAAGAGTGGCATTCAACTGCACACAGGGATTGTTG |
| SATB2-R389C-R | CAACAATCCCTGTGTGCAGTTGAATGCCACTCTTG |
| SATB2-R389L-F | AAGAGTGGCATTCAACCTCACACAGGGATTGTTGT |
| SATB2-R389L-R | ACAACAATCCCTGTGTGAGGTTGAATGCCACTCTT |
| SATB2-T390I-F | AGAGTGGCATTCAACCGCATACAGGGATTGTTGT |
| SATB2-T390I-R | ACAACAATCCCTGTATGCGGTTGAATGCCACTCT |
| SATB2-Q391R-F | GCATTCAACCGCACACGGGGATTGTTGTCTGAG |
| SATB2-Q391R-R | CTCAGACAACAATCCCCGTGTGCGGTTGAATGC |
| SATB2-G392R-F | CATTCAACCGCACACAGCGATTGTTGTCTGAGATT |
| SATB2-G392R-R | AATCTCAGACAACAATCGCTGTGTGCGGTTGAATG |
| SATB2-G392E-F | CATTCAACCGCACACAGGAATTGTTGTCTGAGATTCT |
| SATB2-G392E-R | AGAATCTCAGACAACAATTCCTGTGTGCGGTTGAATG |
| SATB2-G392V-F | CATTCAACCGCACACAGGTATTGTTGTCTGAGATTCT |
| SATB2-G392V-R | AGAATCTCAGACAACAATACCTGTGTGCGGTTGAATG |
| SATB2-L394S-F | CGCACACAGGGATTGTCGTCTGAGATTCTGCGT |
| SATB2-L394S-R | ACGCAGAATCTCAGACGACAATCCCTGTGTGCG |
| SATB2-E396Q-F | TACGCAGAATCTGAGACAACAATCCCTGTGTGCGG |
| SATB2-E396Q-R | CCGCACACAGGGATTGTTGTCTCAGATTCTGCGTA |
| SATB2-R399H-F | GTTGTCTGAGATTCTGCATAAGGAAGAAGACCCTC |
| SATB2-R399H-R | GAGGGTCTTCTTCCTTATGCAGAATCTCAGACAAC |
| SATB2-R399P-F | GTTGTCTGAGATTCTGCCTAAGGAAGAAGACCCTC |
| SATB2-R399P-R | GAGGGTCTTCTTCCTTAGGCAGAATCTCAGACAAC |
| SATB2-R399L-F | GTTGTCTGAGATTCTGCTTAAGGAAGAAGACCCTC |
| SATB2-R399L-R | GAGGGTCTTCTTCCTTAAGCAGAATCTCAGACAAC |
| SATB2-E401K-F | CTGAGATTCTGCGTAAGAAAGAAGACCCTCGGACA |
| SATB2-E401K-R | TGTCCGAGGGTCTTCTTTCTTACGCAGAATCTCAG |
| SATB2-E402K-F | CTGAGATTCTGCGTAAGGAAAAAGACCCTCGGACA |
| SATB2-E402K-R | TGTCCGAGGGTCTTTTTCCTTACGCAGAATCTCAG |
| SATB2-Q409R-F | CCTCGGACAGCCTCTCGGTCTCTTCTAGTAAACC |
| SATB2-Q409R-R | GGTTTACTAGAAGAGACCGAGAGGCTGTCCGAGG |
| SATB2-M418R-F | CTAGTAAACCTGAGGGCCAGGCAGAATTTCCTCAATC |
| SATB2-M418R-R | GATTGAGGAAATTCTGCCTGGCCCTCAGGTTTACTAG |
| SATB2-R429Q-F | CTGCCAGAAGTGGAGCAAGATCGCATCTACCAG |
| SATB2-R429Q-R | CTGGTAGATGCGATCTTGCTCCACTTCTGGCAG |
| SATB2-R431H-F | GAAGTGGAGCGAGATCACATCTACCAGGATGAG |
| SATB2-R431H-R | CTCATCCTGGTAGATGTGATCTCGCTCCACTTC |
| SATB2-E436V-F | CATCTACCAGGATGTGAGGGAGCGGAGCA |
| SATB2-E436V-R | TGCTCCGCTCCCTCACATCCTGGTAGATG |
| SATB2-Q514R-F | TGGCTGCAAATAAAAGTCGGGGCTGGCTGTGTGA |
| SATB2-Q514R-R | TCACACAGCCAGCCCCGACTTTTATTTGCAGCCA |
| SATB2-G515S-F | GCTGCAAATAAAAGTCAGAGCTGGCTGTGTGAACT |
| SATB2-G515S-R | AGTTCACACAGCCAGCTCTGACTTTTATTTGCAGC |
| SATB2-C518W-F | GGGCTGGCTGTGGGAACTGCTCCGC |
| SATB2-C518W-R | GCGGAGCAGTTCCCACAGCCAGCCC |
| SATB2-R522C-F | CTGTGTGAACTGCTCTGCTGGAAGGAGAACC |
| SATB2-R522C-R | GGTTCTCCTTCCAGCAGAGCAGTTCACACAG |
| SATB2-L545P-F | CATCCGTCGCTTCCCGAACCTTCCCCAGC |
| SATB2-L545P-R | GCTGGGGAAGGTTCGGGAAGCGACGGATG |
| SATB2-V554I-F | CCCCAGCATGAGAGGGATATCATCTATGAGGAGG |
| SATB2-V554I-R | CCTCCTCATAGATGATATCCCTCTCATGCTGGGG |

| | |
|---|---|
| SATB2-E566K-F | AGGCATCACCACAGCAAACGCATGCAACACG |
| SATB2-E566K-R | CGTGTTGCATGCGTTTGCTGTGGTGATGCCT |
| SATB2-D635Y-F | GGGATCCTCCAAAGCTTTATTCATTATGTAGGCCTGTA |
| SATB2-D635Y-R | TACAGGCCTACATAATGAATAAAGCTTTGGAGGATCCC |
| SATB2-H646T-F | GACCAGGAAGCCATCTACACTCTTTCGGCTC |
| SATB2-H646T-R | GAGCCGAAAGAGTGTAGATGGCTTCCTGGTC |
| SATB2-S649L-F | AGGAAGCCATCCACACTCTTTTGGCTCAGCTGG |
| SATB2-S649L-R | CCAGCTGAGCCAAAAGAGTGTGGATGGCTTCCT |
| SATB2-P655L-F | GCTCAGCTGGATCTCCTCAAACACACCATCATC |
| SATB2-P655L-R | GATGATGGTGTGTTTGAGGAGATCCAGCTGAGC |
